# Comparative analysis of wavelength-specific UV stress granule formation

**DOI:** 10.64898/2026.03.15.711948

**Authors:** Alexandra J. Cabral, Natalie G. Farny

**Affiliations:** Department of Biology and Biotechnology, Worcester Polytechnic Institute, Worcester, MA, USA; Program in Bioinformatics and Computational Biology, Worcester Polytechnic Institute, Worcester, MA, USA

**Keywords:** Stress granules, ultraviolet radiation (UV), keratinocyte, translational control

## Abstract

Stress Granules (SGs) are cytoplasmic biomolecular condensates that form in response to a variety of stress conditions, though their function remains unclear. “Canonical” SGs – caused by stressors like sodium arsenite – are dynamic and cytoprotective, allowing cells to evade cell death during periods of stress. Ultraviolet (UV) irradiation is known to elicit a “non-canonical” SG subtype, lacking canonical SG components such as eukaryotic initiation factor 3 and polyadenylated mRNAs. The exact function of UV SGs, and the mechanisms driving their formation, remain unknown. Here we report the findings of a comparative analysis of UVA, UVB and UVC exposures on SG formation in three cell types: osteosarcoma (U2OS), keratinocytes (HaCaT), and mouse embryonic fibroblasts (MEF). We observed that SG formation in response to UV is highly cell type dependent. UVB and UVC induce robust SG formation in U2OS cells. However, only UVC exposure induced modest SG formation in MEFs, and none of the wavelengths caused SGs in HaCaT. While UVC-induced SGs in U2OS cells appear to be cell cycle dependent and specific to G1, UVB induced SG formation regardless of cell cycle stage. We tested the hypothesis that oxidative stress triggered by UV may be driving UV SG formation, and that keratin may buffer this effect, by overexpressing keratin in U2OS. Interestingly, we found that keratin and antioxidant treatment efficiently suppressed arsenite-induced SGs but had no effect on UV SGs. Our work confirms that UV SG formation is cell type specific and is not driven by oxidative stress.

## INTRODUCTION

Ultraviolet (UV) light has multiple damaging effects at the cellular level. UV irradiation causes damage to mammalian cells primarily by direct DNA damage, which induces mutations and other photoproducts that can accumulate and drive various cancers and diseases of the skin^1–3^. Along with DNA damage, UV light also induces RNA damage and increases the production of reactive oxygen species (ROS), leading to ROS-mediated cellular damage^4,5^. The UV light spectrum is divided into three wavelength ranges – UVA (315-400nm), UVB (280-315nm), and UVC (100-280nm). While all UV wavelengths cause both DNA damage and ROS to some extent, UVC and UVB generate more direct DNA damage through cyclobutene pyrimidine dimers (CPDs) and pyrimidine 6-4 pyrimidone photoproducts (6-4PPs), whereas UVA generates fewer direct photoproducts and greater ROS^6–10^. Still, the wavelength-specific effects of UV exposure on various cellular processes remain unclear.

Canonical stress granules (SGs) form in response to stress-induced translational repression downstream of the integrated stress response (ISR), and re-program cellular signaling cascades by harboring specific proteins and mRNAs to promote cell survival during brief stress periods^11–17^. Stress granules (SGs) were first observed to form in mammalian cells in 1999 in response to sodium arsenite (As^III^), heat shock and UVC exposure^18^. Canonical SGs have since been shown to be dynamic, cytoprotective, anti-apoptotic, and to disassemble upon the end of stress via the autophagy pathway^15,19–23^. Alternatively, non-canonical SGs have recently been observed and form in response to stressors such as nitric oxide, sodium selenite, chronic nutrient starvation and UV irradiation^24–33^. Non-canonical SGs were originally categorized based on their lack of canonical SG component eukaryotic initiation factor 3 (eIF3)^24^. These non-canonical SGs are reported to be less dynamic and cytotoxic, making them more akin to aggregation seen in disease states^24,30,31^.

Until recently, there were few molecular details known about the formation of UV-induced SGs. UV SGs are driven by G3BP-dependent mechanisms^28^, however they lack both poly(A) RNA and eIF3^34^. They form in only a minor percentage of an asynchronous cell population, and early reports suggested that they may be cell cycle-specific in their formation, forming exclusively in the G1 phase^25,27^. Then, a recent report suggested UV SGs form in cells in G1 phase as a product of UV-induced RNA damage occurring in G2/M phases, as a mechanism to protect daughter cells against UV-induced RNA damage inherited from irradiated parent cells^5^. Therefore, these UV-induced SGs are reported to form as a protective measure, to ensure daughter cell survival, thereby contradicting hypotheses that non-canonical SG subtypes are inherently cytotoxic^5^.

While the pool of UV-induced SG literature is expanding, most prior studies were conducted in cell types that are not physiologically relevant to UV exposure. Keratinocytes make up the outermost protective layer of the skin and are therefore the primary cell type exposed directly to UV irradiation. Our prior work showed that the keratinocyte cell line HaCaT is highly resistant to UVC-induced SG formation^34^. Zhou et al.^5^ included several skin cell lines (N/TERT-1, A431 and HaCaT) within their study to investigate the RNA damaging effects of UV irradiation, and observed UV SG formation primarily in response to UVA after pre-treatment with uridine analog 4-thiouridine (4sU)^5^, which is selectively incorporated into RNA and damaged by UVA^35^. Because UVA induces negligible amounts of DNA damage, these authors concluded that DNA damage is not the primary mechanism contributing to UV-induced SG formation^5^, consistent with a prior report^25^. However, the extent to which UVB or UVC induces SGs within keratinocytes has not been thoroughly quantified, and the underlying mechanism of the relative UV-resistance to SG formation in keratinocyte-derived cells remains unknown.

Reported herein, we show that UVC is the only UV type that induces SGs at a low dose (15mJ/cm^2^) in U2OS (osteosarcoma) cells, and to a much lesser extent in mouse embryonic fibroblasts (MEFs) and HaCaT cells. UVB at higher doses (150mJ/cm^2^) induces robust SGs in U2OS cells, but not in HaCaT cells, and UVA alone does not induce SGs in any cell type. We show that UVB and UVC-induced SGs are similar in terms of key protein composition. Interestingly, we observe that while UVC-induced SGs only form in cells in the G1 phase of the cell cycle, UVB-induced SGs form in ∼60% of U2OS cells regardless of cell cycle stage. It is well established that UV induces oxidative stress^4,6,36^, and that oxidative stress induces SG formation^37^. Therefore, we hypothesized that oxidative stress caused by UVB irradiation could be driving the robust SG formation we observe in asynchronously growing U2OS cells. Further, we hypothesized that the UV SG suppression affect in HaCaT cells may be due to the antioxidant properties of keratin proteins^38–40^. To this end we show that the expression of the keratin pair K8/K18 in U2OS cells does not suppress UVB-induced SGs. Interestingly, we find that keratin expression suppresses canonical, oxidative As^III^-induced SGs in U2OS cells. We propose that keratin is combatting the oxidative stress induced by As^III^, and that any oxidative stress induced by UVB is insufficient to drive the robust SG formation we observed in U2OS cells.

## RESULTS

### UVC induces SGs at low doses while UVB induces SGs only at higher doses

While UVC has been reported to induce cytoplasmic SGs, fewer studies have focused on understanding the effects of UVA and UVB on SG formation^25–28^. We show that at low doses (15 mJ/cm^2^) only UVC induces SG formation in U2OS cells (Fig. 1, Fig. 2B) and to a lesser extent in HaCaT (Fig. 2A-B) and MEF cells (Fig. 2B, Supplementary Fig. S2). At higher doses (150 mJ/cm^2^) UVB induces robust SG formation in U2OS cells; however, UVC SGs do not increase significantly relative to the lower 15 mJ/cm^2^ dose (Fig. 2C). Interestingly, we observe UVB at 150mJ/cm^2^ induces ∼60% SG formation in U2OS cells while previous studies have reported ∼30% SG formation in response to UVB in HeLa cells^5^.

**Figure 1.**
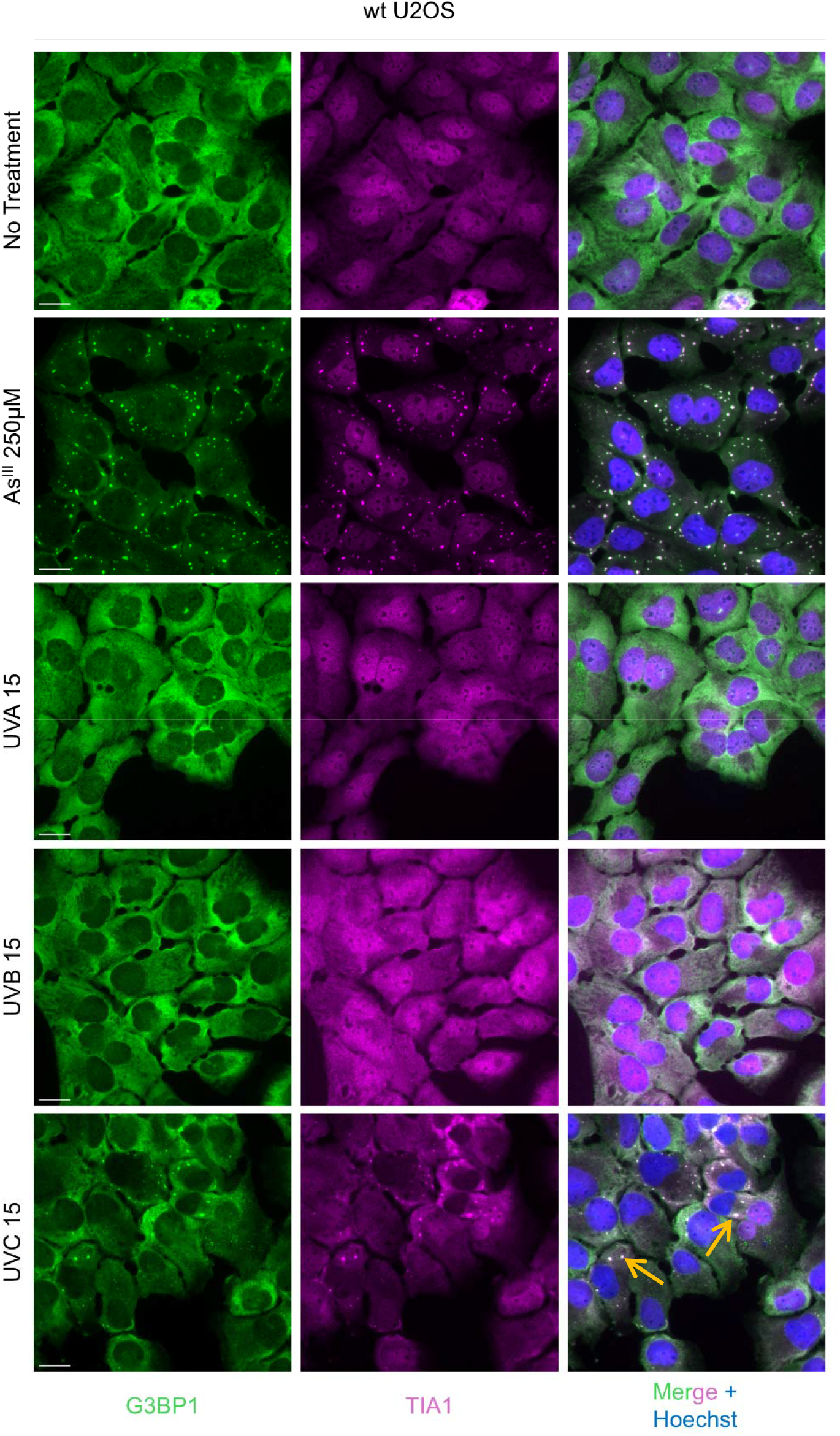
UVC induces SGs in U2OS cells at low doses. Representative images of SG formation by G3BP1 and TIA1 signals in wt U2OS cells treated as indicated. Cells were either untreated, treated with 250μM As^III^ for 1h, or exposed to 15mJ/cm^2^ of UV. UV samples were harvested and processed for imaging 4 hours after exposure. Yellow arrows indicate SGs. Scale bar = 20μm.

**Figure 2.**
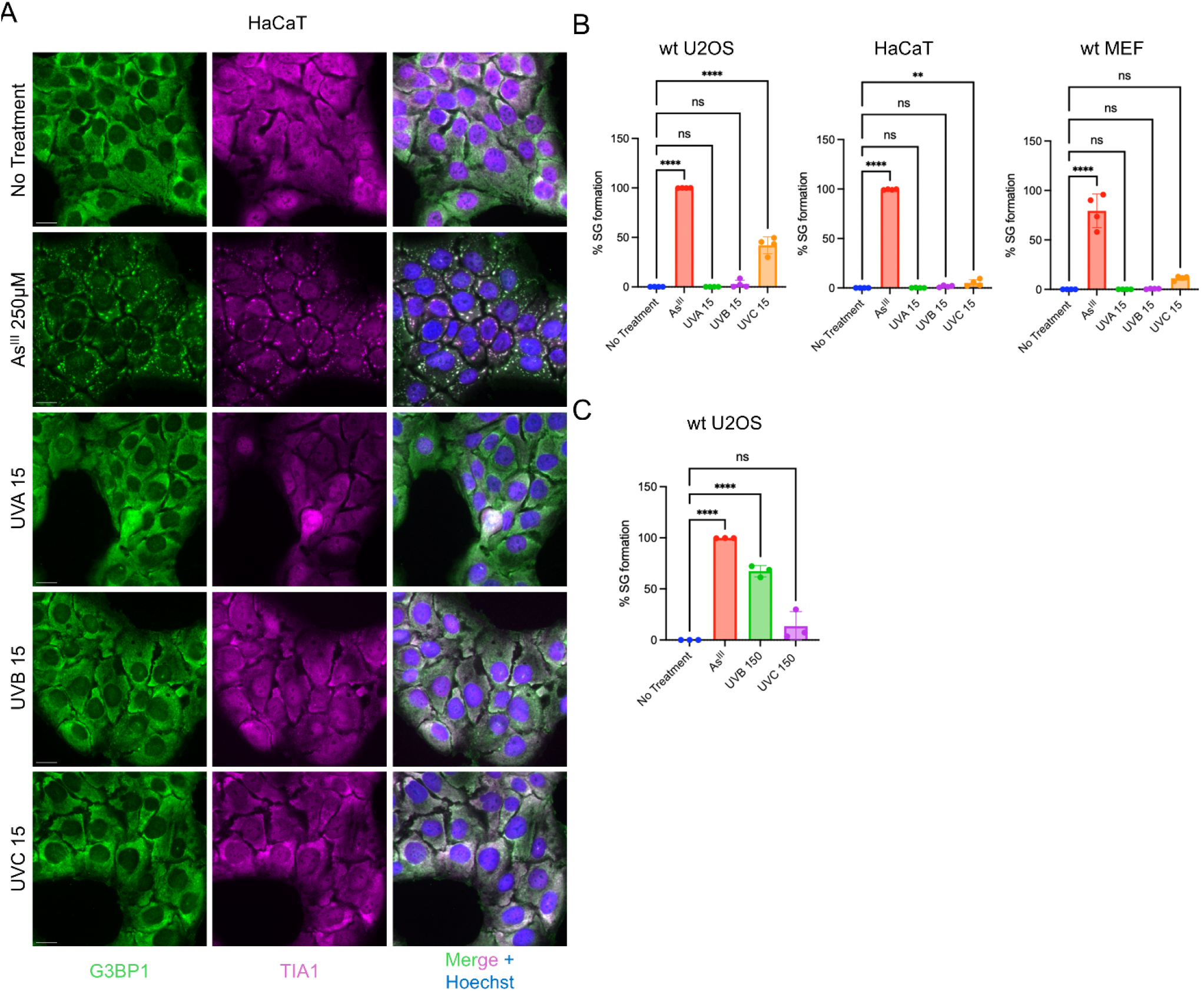
Low doses of UV do not robustly induce SGs in HaCaT or MEF cells while high doses of UVB induce SG in U2OS cells. (A) Representative images of SG formation by G3BP1 and TIA1 signals in HaCaT cells treated as indicated. Cells were either untreated, treated with 250μM As^III^ for 1h, or exposed to 15mJ/cm^2^ of UV. UV samples were harvested and processed for imaging 4 hours after exposure. (B) Quantification of SG formation by G3BP1 and TIA1 signals in wt U2OS, HaCaT, and wt MEF cells treated as indicated. Cells were either untreated, treated with 250μM As^III^ for 1h, or exposed to 15mJ/cm^2^ of UV. UV samples were harvested and processed for imaging 4 hours after exposure. At least 250 cells and 3 fields of view were scored to calculate percent SG formation. *n*=3; error bars are ±SD; ns P > 0.05, ** P ≤ 0.01, **** P ≤ 0.0001 by one-way ANOVA and Dunnett’s post-hoc test. (C) Quantification of SG formation by G3BP1 and TIA1 signals in wt U2OS cells treated as indicated. Cells were either untreated, treated with 250μM As^III^ for 1h, or exposed to 150mJ/cm^2^ of UV. UV samples were harvested and processed for imaging 4 hours after exposure. At least 250 cells and 3 fields of view were scored to calculate percent SG formation. *n*=3; error bars are ±SD; ns P > 0.05, **** P ≤ 0.0001 by one-way ANOVA and Dunnett’s post-hoc test. Scale bar = 20μm.

It is well established that UV irradiation activates cellular stress response pathways as a result of UV-induced DNA and ROS-mediated damage^1,2,4,6^. In addition to triggering these pathways, UV exposure also decreases cell viability^41–43^. To confirm that our UV treatments induced the expected damage responses and decreased cell viability, we performed preliminary western blot analyses for markers of UV damage (PARP cleavage and ERK phosphorylation) and MTT assays to measure cell viability (Supplementary Fig. S3). Collectively, our preliminary investigations suggest that UV damage is occurring in the treated cells as expected, and therefore the lack of SG formation under some contexts is not likely to be due to a lack of UV damage.

### UVB and UVC-induced SGs are compositionally similar

Canonical SGs are reported to sequester a variety of different proteins, and the localization of certain proteins is used to determine which downstream stress response pathways may be triggered to combat a specific stressor. Canonically, eIF3 is a major component of SGs as well as the small ribosomal subunit and translationally stalled mRNAs^14,44^. These proteins and mRNAs are recruited due to their roles in the eukaryotic pre-initiation complex required for translation initiation. During stress bulk translation is inhibited which results in the accumulation of stalled pre-initiation complexes in the cytoplasm that become major components of SGs^11^. However, UV SGs have been dubbed non-canonical, and this conclusion was drawn initially due to the lack of canonical SG markers such as eukaryotic initiation factor 3 (eIF3) and poly (A) RNA^24^. Recently, UV SGs were reported to recruit the double stranded RNA helicase DHX9, a protein that is not enriched in canonical SGs^5^. These authors also uncovered that damaged intron RNA is recruited to these granules^5^.

Due to the conflicting reports regarding canonical versus non-canonical SG protein composition, as well as conflicting opinions on whether or not UV SGs are cytoprotective or pro-apoptotic, we performed immunofluorescence microscopy to compare UVB-induced SG composition to UVC-induced SG composition, using As^III^-induced SGs as a canonical control. Our findings corroborated recent reports that UVC-induced SGs and UVB-induced SGs recruit DHX9 (Figure 3A). Additionally, consistent with previous findings, we show that pre-initiation complex components eIF3n and ribosomal protein S6 (Rps6) are excluded from UVB and UVC SGs (Figure 3B-C). We also show that mature poly (A) RNA is absent from UVB and UVC induced SGs (Figure 3D). Co-localization plot profiles were used to confirm these observations (Supplementary Fig. S4). Our findings indicate that UVB- and UVC-induced SGs are compositionally similar, suggesting they may have comparable function.

**Figure 3.**
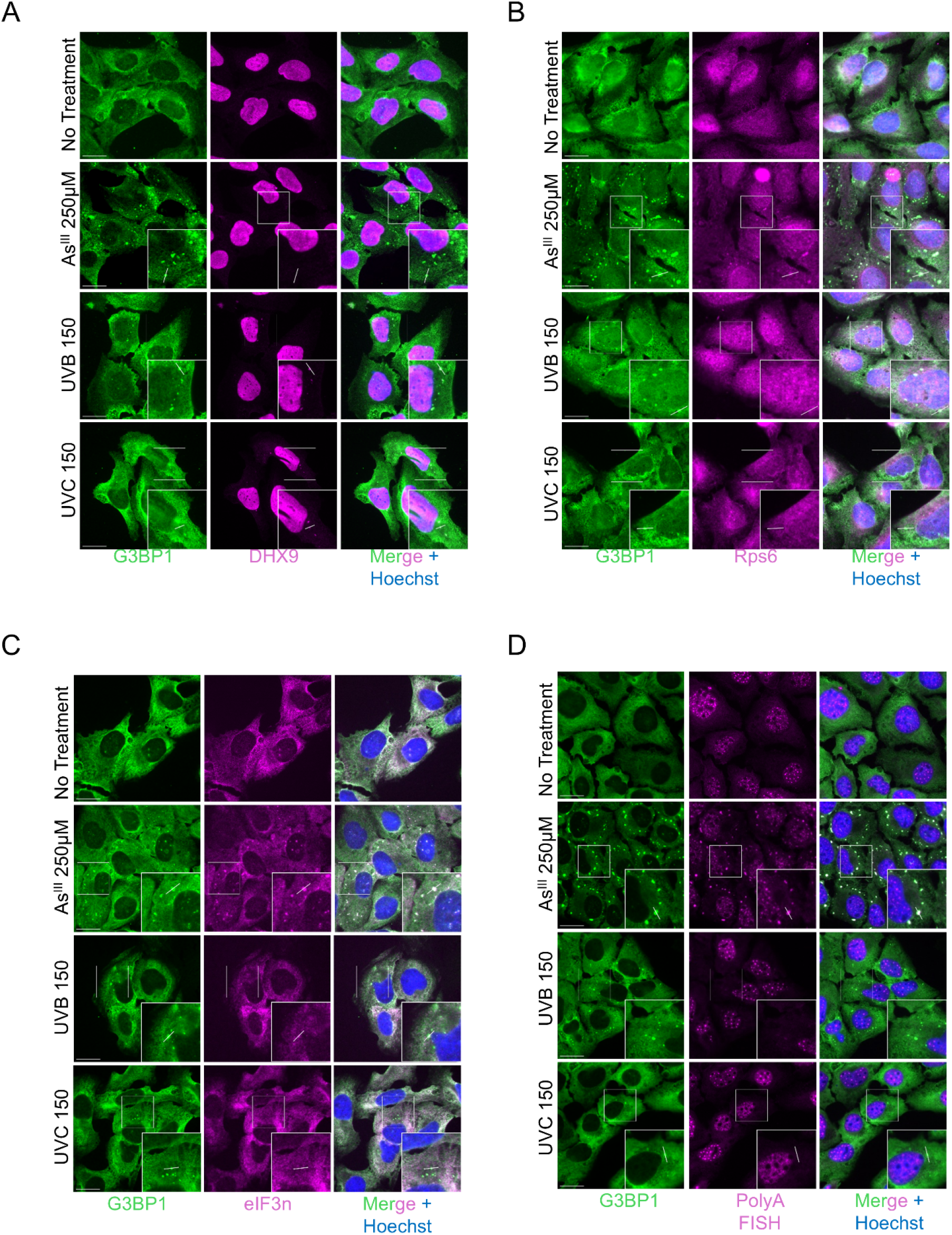
UVB and UVC-induced SGs are compositionally identical. (A-D) Representative images of SGs identified by G3BP1 co-stained with relevant SG markers (A) DHX9, (B) Rps6, (C) eIF3n, (D) Poly(A) fluorescence in situ hybridization (FISH). wt U2OS cells were either untreated, treated with 250μM As^III^ for 1h, or exposed to 150mJ/cm^2^ of UV. UV samples were harvested and processed for imaging 4 hours after exposure. Cells were stained with relevant SG marker antibodies and observed and imaged under 63X magnification. Scale bar = 20μm.

### UVB SGs do not form in a cell-cycle dependent manner in U2OS cells

UV SGs were reported to form in a cell-cycle dependent manner in several prior reports, appearing only in post-mitotic cells in G1 following UVC^25,27^ and UVA+4sU^5^ exposures. However, we observed that in asynchronously growing U2OS cells, UVB induced robust (∼60%) SG formation (Figure 2C). Given the high percentage of SG-positive cells under UVB exposure, we thought it unlikely that all cells that formed SGs were in the same cell cycle phase. Therefore, we hypothesized that UVB-induced SGs are not cell cycle dependent like their UVC and UVA+4sU counterparts. To test this hypothesis, we performed immunofluorescence microscopy to visualize UV SGs and the cycle protein geminin. Geminin is a central regulator in the cell cycle and is localized to the nucleus during S, G2 and M phases. Therefore, if UVB-induced SGs form in the same manner as UVC and UVA+4sU-induced SGs we would expect to observe them exclusively in cells where nuclear geminin is absent. Interestingly, we observe that UVB-induced SGs form in U2OS cells with and without nuclear geminin, suggesting that they do not form in a cell cycle dependent manner (Fig. 4). This result implies that the mechanisms that drive UVC and UVB-induced SGs in U2OS cells may be distinct. Thus, although DHX9 is recruited to both UVC and UVB-induced SGs, it is unlikely that RNA damage is the only factor contributing to UVB-induced SGs in U2OS cells.

**Figure 4.**
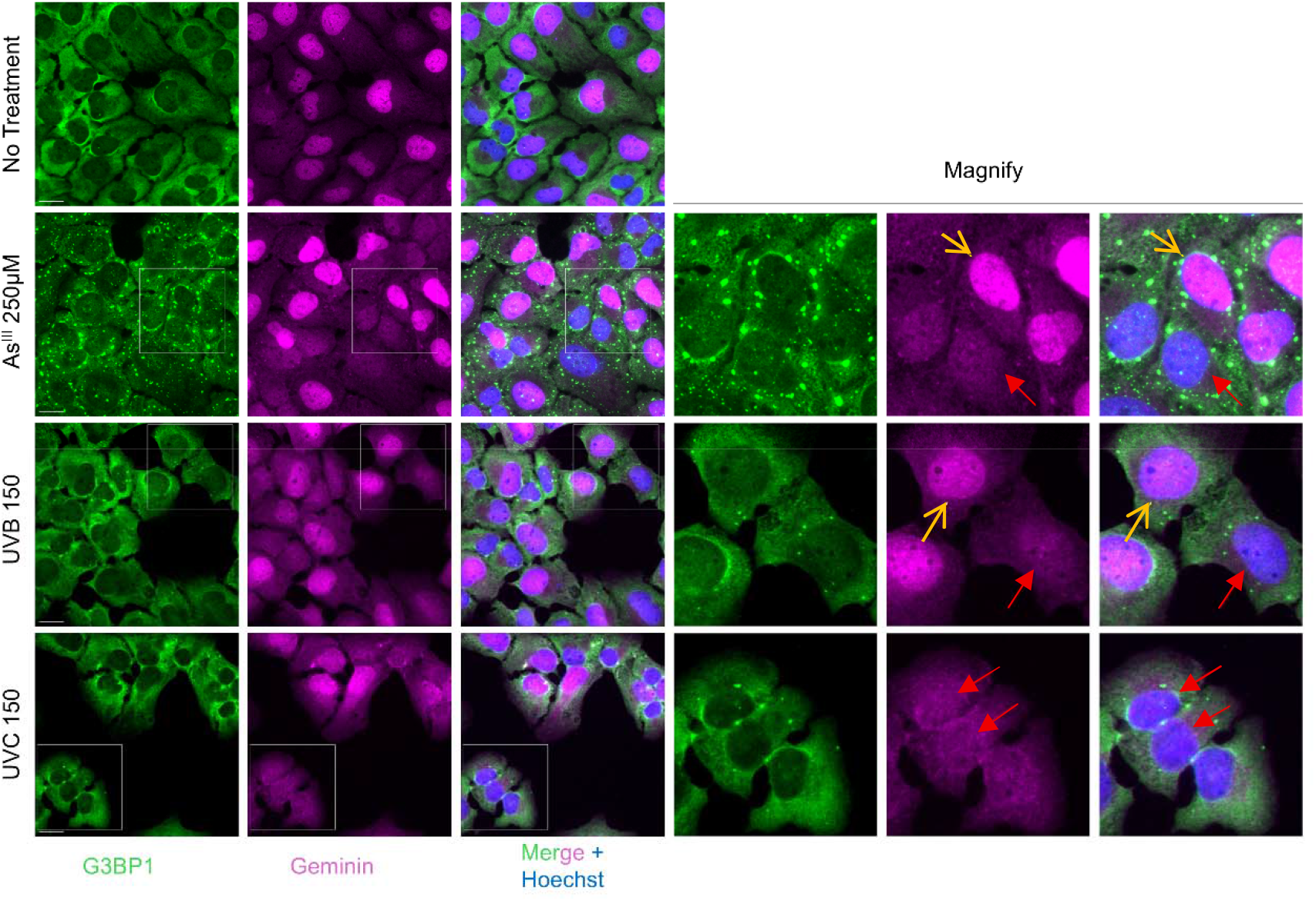
UVB-induced SGs may not be cycle dependent in U2OS cells. Representative images of wt U2OS cells treated as indicated and stained for G3BP1 and Geminin. Cells were either untreated, treated with 250μM As^III^ for 1h, or exposed to 150mJ/cm^2^ of UV. UV samples were harvested and processed for imaging 4 hours after exposure. Yellow arrows indicate cells in S, G2, or M phase. Red arrows indicate cells in G1 phase. Scale bar = 20μm.

### HaCaT cells do not robustly form UV-induced SGs

Much of the available literature investigates UV-induced SGs in physiologically irrelevant cell types^27,28^. Skin cells, and specifically keratinocytes, are the outermost layer of the skin and are therefore our first line of defense against damaging UV irradiation. We observed that HaCaT cells do not robustly form UV SGs in response to 15 mJ/cm^2^ of UVA, UVB, or UVC (Figure 2A-B). Still, it was possible that these doses were insufficient to cause SGs in HaCaTs. To fully probe the ability of HaCaT cells to form UV SGs and to eliminate the possibility that we had not adequately dosed the cells in order to observe UV SGs in HaCaTs, we exposed HaCaTs to a high-dose (150mJ/cm^2^) of UVB and UVC. Our results show that HaCaT cells do not form UV SGs in response to higher doses (150mJ/cm^2^) of UVC, and only form in a few isolated cells (2-6% of cells) in response to UVB (Supplementary Figure S5D). The few UVB-induced SGs that form in HaCaT cells recruit G3BP1, TIA-1, and DHX9 (Supplementary Fig. S5A-B) and, conversely, these SGs are not enriched in Rps6 (Supplementary Fig. S5C). Furthermore, we find that even when HaCaT cells are pre-treated with 4sU there is no significant SG formation in response to UVA at 150mJ/cm^2^ (Supplementary Fig. S6). Thus, although we report minimal UV-induced SG formation in HaCaT cells (Supplementary Figs. S5 and S6), we do observe that the small population of cells that form SGs do so in the same manner described by Zhou et al^5^. Further, in HaCaT cells we observe these SGs exclusively in cells without nuclear geminin signal (Supplementary Fig. S7), suggesting that they form in the cell cycle-dependent manner previously described.

While we observed some SG formation in response to UVB and UVA+4sU (Supplementary Fig. S5 and S6), the results shown here, along with our previous report^34^, suggest that HaCaT cells have an intrinsic protective mechanism to suppress UV-induced SG formation. To examine whether UVA+4sU would cause SG formation in our hands, we applied this treatment to U2OS cells. We observed that U2OS cells form a small but significant (∼3%) amount of UVA+4sU-induced SGs (Fig. 5A-B). These findings, coupled with our observation that UVB-induced SGs are not cell cycle dependent in U2OS cells, imply that the mechanisms involved in UVB-induced SG formation may be cell-type specific.

**Figure 5.**
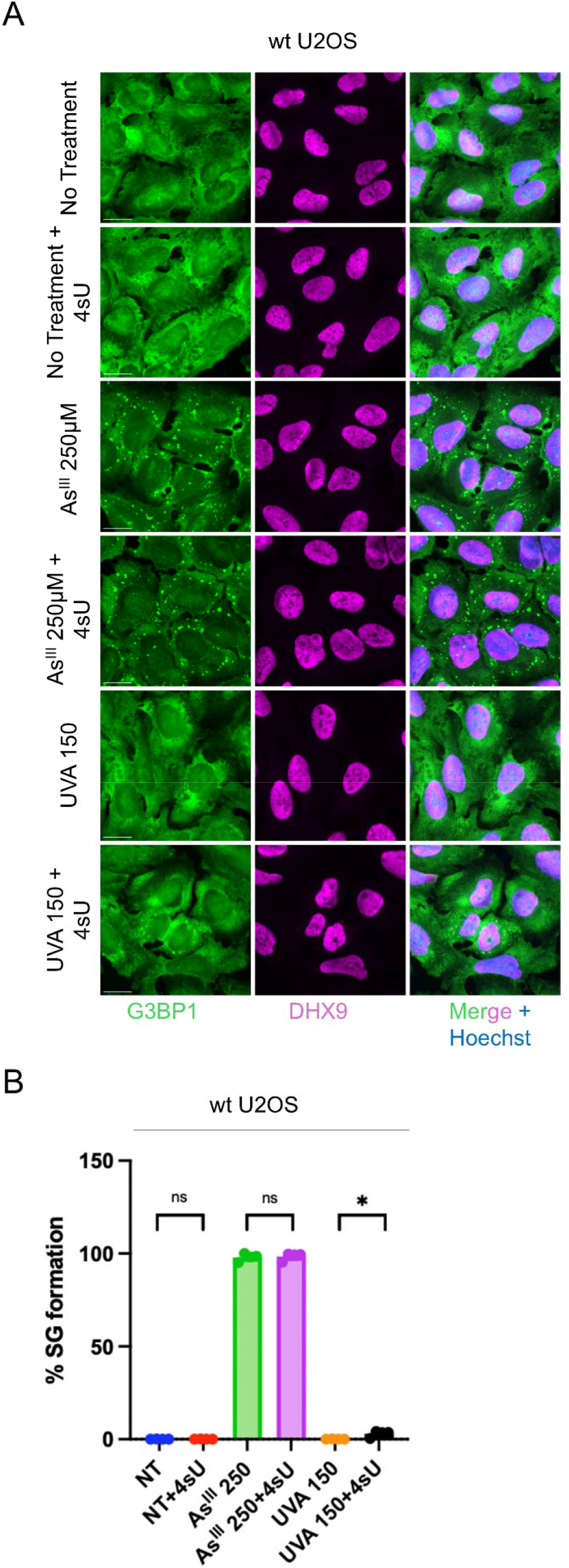
Minimal SGs form in U2OS cells in response to UVA + 4sU. (A-B) Representative images (A) and quantification (B) of SG formation in wt U2OS cells after treatment as indicated. Cells were either untreated or pre-treated with 4sU for 1h. Following pretreatment cells were either untreated, treated with 250μM As^III^ for 1h, or treated with UVA. UV samples were harvested and processed for imaging 8 hours after exposure. At least 250 cells and 3 fields of view were scored to calculate percent SG formation. *n*=4; error bars are ±SD; ns P > 0.05, * P≤0.05 by unpaired *t* test. Scale bar = 20μm.

### Keratin expression suppresses As^III^-induced SGs but not UV-induced SGs

Thus far, our results suggest that SGs that from in response to UVB in U2OS cells do so in a manner distinct from those that form in response to UVC. This finding, coupled with our observation that HaCaT cells possess an intrinsic protective mechanism that suppresses SG formation in response to UV, led us to hypothesize that UV-induced oxidative stress may be contributing to UVB-induced SG formation in U2OS cells. Prior reports state that DNA damage alone is insufficient to induce SG assembly^5^, however it is well established that oxidative stress favors SG formation^37^. Therefore, we hypothesized that UVB-induced oxidative stress may be driving the increased SG formation in response to UVB we observe in U2OS cells that are not exclusively in the G1 phase.

Additionally, we speculated that the SG suppression we observe in HaCaT cells is a result of oxidative stress buffering by the expression of keratin intermediate filaments in the cytoplasm. Keratins are a family of cytoskeletal proteins that are subdivided into Type I and Type II keratins. Type I and Type II keratins are interdependent; one of each type is required for proper intermediate filament formation^45^. Keratins have been reported to have antioxidant properties^38–40^ and therefore we hypothesized that the expression of keratin intermediate filaments in U2OS cells could suppress UVB-induced SG formation by buffering oxidative stress.

To determine whether keratins can inhibit UV SG formation, we overexpressed the keratin pair K18 (Type I) and K8 (Type II) in U2OS cells, which endogenously express keratin only at a very low level (Fig. 6A-B). In the K8/K18 transfected cells, we did not observe any suppression effect in UVB or UVC-induced SG formation (Figure 6C). Interestingly, we did observe the suppression of As^III^-induced SGs in K8/K18-transfected cells; this finding supports the evidence that keratins have antioxidant properties and confirms those properties to be robust enough to buffer oxidative stress and suppress oxidation-induced SG formation. Although we did not observe suppression in UVB or UVC SG formation in response to K8/K18 expression, further analysis was necessary to confirm that oxidative stress is not contributing to UVB-induced SG formation.

**Figure 6.**
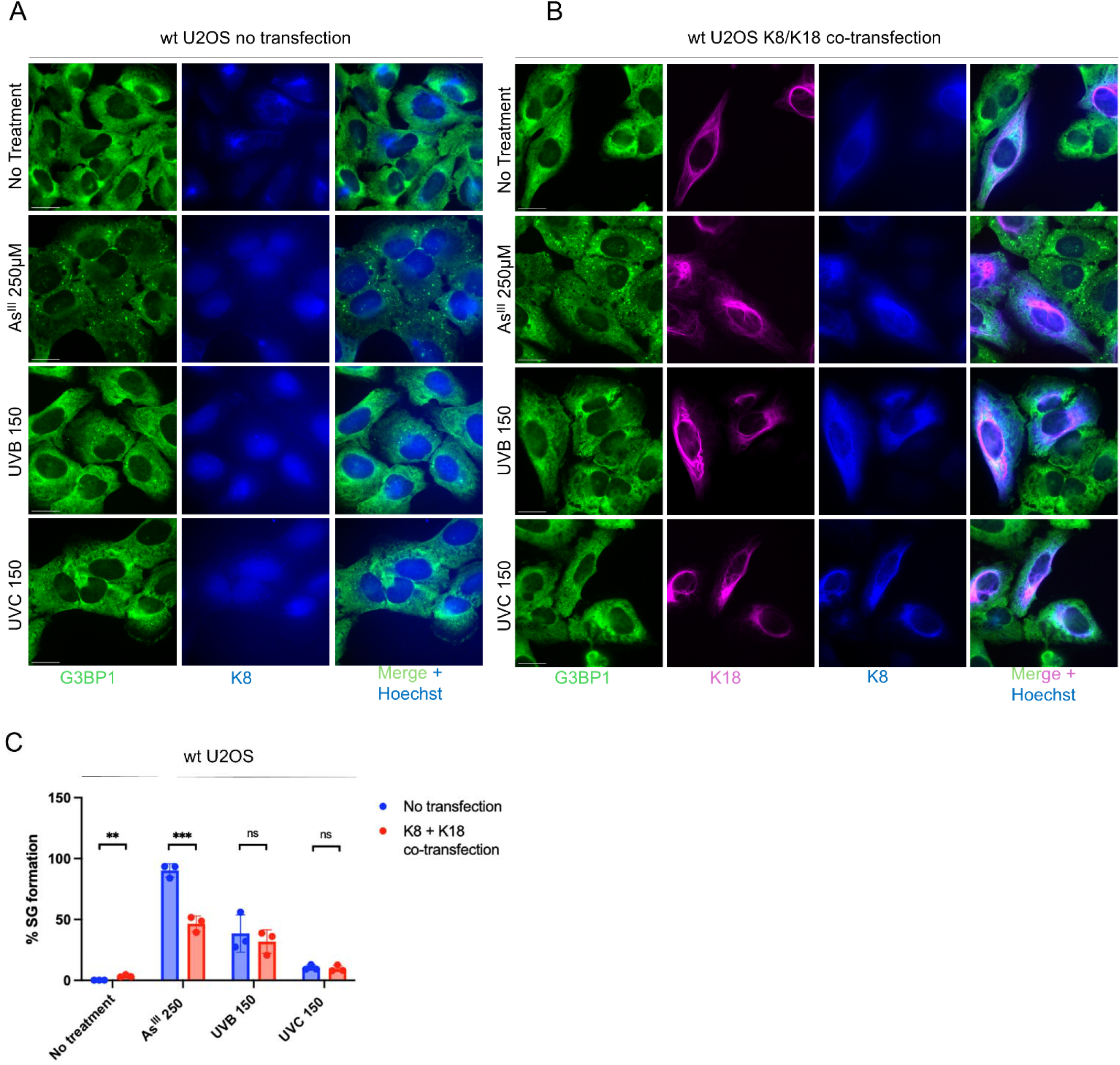
K8 and K18 overexpression suppresses As^III^-induced SG formation but not UV-induced SG formation. (A) Representative images of wt U2OS cells stained for G3BP1 and keratin 8 treated as indicated. Cells were either untreated, treated with 250μM As^III^ for 1h, or exposed to 150mJ/cm^2^ of UV. UV samples were harvested and processed for imaging 4 hours after exposure. (B) Representative images of wt U2OS cells co-transfected with keratin 8 and keratin18_mCherry. Cells were either untreated, treated with 250μM As^III^ for 1h, or exposed to 150mJ/cm^2^ of UV. UV samples were harvested and processed for imaging 4 hours after exposure. (C) Quantification of SG formation in untransfected and co-transfected cells by G3BP1 staining. Cells were treated as indicated. At least 200 cells and 3 fields of view were scored to calculate percent SG formation. *n*=3; error bars are ±SD; ns P > 0.05, ** P ≤ 0.01, *** P ≤ 0.001 by unpaired *t*-test. Scale bar = 20μm.

### UVB-induced SGs are not robustly driven by oxidative stress

N-acetylcysteine (NAC) is a well-characterized antioxidant and has been previously reported to suppress oxidative stress-induced SG formation, including that triggered by arsenite and hydrogen peroxide^33,46,47^. To confirm that oxidative stress is not robustly contributing to UVB-induced SG formation in U2OS cells, we co-treated cells with NAC and As^III^, UVB, or UVC. Consistent with prior reports^33,46,47^, NAC suppressed As^III^-induced SGs; however, NAC co-treatment did not suppress UVB- or UVC-induced SG formation (Fig. 7A-B). Together, these findings indicate that UVB-induced SGs in U2OS are not being driven by oxidative stress-dependent mechanisms.

**Figure 7.**
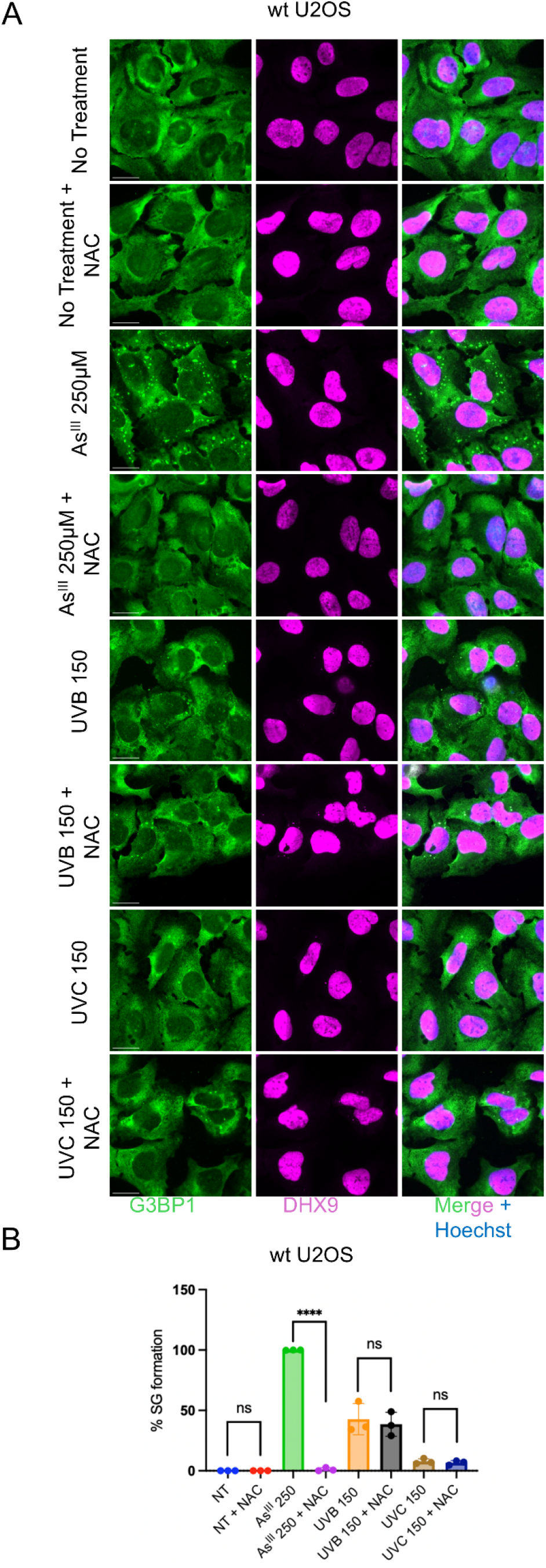
NAC suppresses As^III^-induced SG formation but not UV-induced SG formation. (A-B) Representative images (A) and quantification of SG formation in U2OS cells after treated as indicated. Cells were either untreated, treated with NAC (10mM) for 1h, treated with 250μM As^III^ for 1h, co-treated with NAC and 250μM As^III^ for 1h, treated with UV, or co-treated with UV and NAC. UV samples were harvested and processed for imaging 4 hours after exposure. At least 250 cells and 3 fields of view were scored to calculate percent SG formation. *n*=3; error bars are ±SD; ns P > 0.05, ** P ≤ 0.01, **** P ≤ 0.0001 by unpaired *t*-test.

## DISCUSSION

The field of SG biology has vastly expanded since the discovery of SGs in mammalian cells in 1999^18^. This expansion has led to the identification of numerous SG-inducing agents as well as different SG subtypes^24,29^. Recently, non-canonical SGs have been reported and these SGs are hypothesized to form and function distinctly from canonical SGs^24^. UV irradiation is included in the list of non-canonical SG-inducing agents, however, the pool of literature investigating UV-induced SGs is limited^5,25–28,34^. In literature reviews, non-canonical SGs are speculated to be “pro-apoptotic,” cytotoxic, and less dynamic; however, these conclusions are formed based on the available, yet limited, evidence for all reported non-canonical SGs^24^. It is becoming increasingly likely that different stressors result in compositionally and perhaps functionally distinct SG formation and therefore these broad conclusions may not be true of all non-canonical SGs.

Prior to a report published in 2024^5^, no available studies investigated UV SG formation in any skin cell types. Furthermore, few studies have investigated UV SG formation in response to UVA and UVB, the two most physiologically relevant UV types, as they represent daily UV exposures from the sun (UVC is filtered out by the atmosphere). In our study, we report that at 15 mJ/cm^2^, only UVC induces SG formation in U2OS cells, and to a much lesser extent in MEF and HaCaT cells (Figs. 1 & 2). Additionally, we show that at higher doses (150 mJ/cm^2^), UVB induces robust SG formation in U2OS cells (Fig. 2C), though UV SG formation in HaCaT is minimal at all wavelengths. The exposures used in our study, particularly for UVA and UVB, have physiological relevance. For reference, one study of high school students that were in and out of the school building over the course of a summer day recorded exposures of 5 – 15 mJ/cm^2^ per day^48^. Higher exposures are possible with prolonged time outdoors and at higher altitudes. One study estimates the combined accumulated dosage of UVA and UVB required to cause erythema (sunburn) in individuals with fair skin at 68.7 mJ/cm^2^ ^49^. Thus, our results with HaCaT cells suggest that it is unlikely that SGs are forming in human dermal keratinocytes upon normal solar UV exposures, though these doses may be within SG range for other cell types.

Prior studies have reported that UV SGs form exclusively in the G1 phase of the cell cycle^25,27^ and that these SGs form in daughter cells as a consequence of damaged RNA inherited from the irradiated parent cell^5^. Our finding that ∼60% of U2OS cells exposed to UVB (150 mJ/cm^2^) form SGs (Figure 2C), coupled with our observation that UVB SGs form in U2OS cells with or without nuclear geminin staining (Figure 4), suggests that these SGs may form in a cell-cycle independent manner. The findings reported by Zhou et al.^5^—that UV-induced SGs form in the daughter cells of recently divided mother cells as a protective mechanism against damaged parental RNA—do not fully account for our observations. We therefore propose that an additional mechanism may contribute to UVB-induced SG formation in different cell types.

We hypothesized that this additional mechanism could be related to oxidative stress caused by UV. However, we found that was not the case, as buffering of oxidative stress through keratin overexpression or NAC did not alleviate UV SG formation (Figs. 6 and 7). Zhou et al. also reported that approximately 10% of cells that form UV-induced SGs subsequently resolve these granules and proceed through the cell cycle. Thus, it is possible that UVB-induced SGs initially form in recently divided G1-phase cells, and that a subset of cells can continue progressing through the cell cycle without resolving them. Future studies using live-cell imaging will be necessary to determine whether UVB-induced SG formation is truly cell-cycle independent or occurs exclusively in G1 and persists through the cell cycle.

Like all immortalized cell lines, U2OS and HaCaT have defects in cell cycle regulation that enable them to be passaged continuously and divide indefinitely. In future studies, aligning the specific differences in cell cycle regulation between these two cell lines could lead to new hypotheses about whether distinct cell cycle defects could contribute to the difference in SG formation we observe between these cell lines in response to UVB. Genotoxic stress and specifically UV stress has been reported to induce cell cycle arrest in many cell lines; however, it has been reported that U2OS cells do not arrest the cell cycle upon UVC irradiation^50^. In contrast, HaCaT cell have been reported to arrest the cell cycle in response to UVB irradiation^51^. This distinction suggests that there could be a difference in cell cycle or DNA damage regulation in response to different UV types between these cell types. Future experiments should focus on investigating the cell cycle abnormalities between these two cell lines to establish if these fundamental characteristics could contribute to the difference in UVB-induced SG formation. Additionally, transcriptomic and proteomic studies in unstressed and UV treated (UVB and UVC) cells (U2OS and HaCaT) could illuminate the induction of additional (and potentially differential) stress response pathways. Any differential stress responses between these cell lines could elucidate other types of UV-induced damage that may be contributing to robust SG formation in U2OS that we do not observe in HaCaT cells.

Understanding how UV stress promotes the formation of different types of non-canonical SGs could lend insight into other types of aggregation, for example, protein aggregation seen in human disease. The presence of protein aggregation in disease has been extensively studied, and while there are many disease-specific aggregation types, SGs are directly implicated in amyotrophic lateral sclerosis (ALS), Frontotemporal Lobar Degeneration with TDP-43 lesions (FTLD-TDP), cancer, and Alzheimer’s Disease (AD)^52–58^. The knowledge of stress-specific and cell type-specific SG formation contributes to a broader understanding of both the physiological and pathological roles of biomolecular condensation. The acquisition of such knowledge is essential to the development of therapeutic strategies for aggregation diseases in which SGs are directly implicated.

## MATERIALS AND METHODS

### Cell culture and stress treatments

Cultured cell lines U2OS (ATCC HTB-96), wild-type C57-Bl6 MEF (primary MEFs transformed by the 3T3 method, a gift of Joel D. Richter, UMass Chan Medical School) and HaCaT, were maintained at 37ºC in a CO2 incubator (5% CO2) in DMEM (Gibco) supplemented with 10% FBS, 1% penicillin/streptomycin and 1% glutamine (0.2% glutamine for HaCaT cells). For stress treatments, cells were plated into 12-well plates at 1.5×10^5^ cells/well the day prior to treatment on round glass coverslips. On the day of treatment cells were either left untreated, treated with sodium arsenite (As^III^, 250uM, Sigma, S7400) for 1 hour, or irradiated with indicated doses of UVA (uvBeast™ V1, Amazon), UVB (UV Transilluminator), or UVC (CL-1000 Ultraviolet Crosslinker). UVA and UVB sources were measured using a calibrated UV meter (THOR Labs, PM100D) and a UV sensor (THOR Labs, S120VC 200-1100nm) prior to each stress treatment to ensure accurate dosing (Supplementary Fig. S1 and Supplementary Methods). UV-treated samples were washed with 1X PBS and then irradiated with UV as indicated. After UV irradiation pre-conditioned medium was placed back on cells and samples were harvested for analysis 4 hours post irradiation. For 4sU+UVA treatments HaCaT or U2OS cells were pre-treated with 4sU (500ug/mL, Sigma, T4509) for 1 hour, irradiated with UVA and harvested 8 hours post irradiation. For NAC treatments U2OS cells were co-treated with NAC (10mM) and As^III^ (250μM) for 1 hour, or irradiated with UV and treated with NAC (10mM) in pre-conditioned medium for 4 hours before harvesting.

### Immunofluorescence

Cells on glass coverslips were fixed with 4% paraformaldehyde in PBS for 10 minutes at room temperature, permeabilized with methanol for 10 minutes at room temperature, followed by blocking with 5% BSA in PBS for at least 1 hour at room temperature. Cells were incubated with primary antibodies diluted 1:1000 in 5% BSA for 1 hour at room temperature or overnight at 4ºC. The following antibodies were used in this study; G3BP1 (Proteintech® 13057 (rabbit) and 66486 (mouse)), S6 Ribosomal protein (RPS6) (ABclonal A6058), TIA1 (Proteintech® 12133), RACK1 (Proteintech® 66940), TRAF2 (Proteintech® 26846 (rabbit) 67315 (mouse)), Geminin (Proteintech® 10802), DHX9 (Proteintech®17721), cytokeratin 8 (Invitrogen, 06318), and eIF3n (Santa Cruz, 137214). Coverslips were washed 3 times with 1X PBS and then incubated with secondary antibodies and Hoechst 33342 (Thermo Scientific) for 1 hour at room temperature. Coverslips were washed again 3 times with 1X PBS. Coverslips were then mounted onto glass microscope slides using Vinol. Mature poly (A) mRNA was detected using fluorescence in situ hybridization (FISH). Briefly, cells were treated as indicated, fixed with 4% paraformaldehyde in PBS for 10 minutes, permeabilized with 0.1% triton in PBS for 10 minutes, rehydrated with 70% ethanol for 10 minutes and incubated with oligo-dT_40_ probe (1:200, IDT DNA) overnight at 37ºC. Cells were washed 3 times with 2X saline sodium citrate (SSC) followed by additional antibody staining as described above.

### Microscopy and stress granule quantification

Fluorescence microscopy was performed using a Zeiss Axio Observer A1 inverted Fluorescence Microscope equipped with X-cite 120LED Boost High-Power LED illumination System. Image acquisition was done under 63X magnification with a ThorLabs Kiralux 2.3 MP Monochrome CMOS Camera (ThorLabs CS235MU), and images were processed using ImageJ. SG formation was quantified by manually scoring at least 250 cells and 3 fields of view under 40X magnification. For transfection experiments at least 200 cells were scored. Cells were considered positive for SG formation if they were observed to have at least 2 cytoplasmic SGs.

### Transient transfection

U2OS cells were plated onto glass coverslips at 1.5×10^5^ and the following day transfected with a plasmid containing Keratin 18, a plasmid containing Keratin 8, or co-transfected with both plasmids. Keratin 8 (pcDNA3) was a gift from M Bishr Omary (Addgene plasmid # 18063; http://n2t.net/addgene:18063; RRID: Addgene_18063)^59^ and mCherry-Keratin-17 was a gift from Michael Davidson (Addgene plasmid # 55065; http://n2t.net/addgene:55065; RRID:Addgene_55065). Transfections were conducted using Lipofectamine™ 3000 Transfection Reagent (Thermo Scientific™ L3000001) according to manufacturer’s guidelines. Following transfections cells were treated as indicated and prepared for immunofluorescence microscopy.

### Quantification and statistical analysis

Statistical analysis of an unpaired *t*-test was performed where indicated and statistical significance was considered where P≤0.05. *n* indicates the number of independent biological replicate experiments and within each experiment at least two technical replicates were averaged. Statistical analysis of a one-way ANOVA was performed where indicated followed by Dunnett’s multiple comparisons test. Statistical significance was considered when adjusted P≤0.05. Statistical analysis was performed with GraphPad Prism (Version 10.4.1).

## ACKNOWLEDGEMENTS

We thank the Titova Laboratory at Worcester Polytechnic Institute for assistance with UV meter calibration, and we are especially grateful to Kateryna Friedman for sharing her expertise. We thank Joel D. Richter (UMass Chan Medical School) for MEFs and Lou Roberts (Worcester Polytechnic Institute) for HaCaT cells. N.G.F. is supported by NIH R03AG077140 and R15GM157697.

## AUTHOR CONTRIBUTIONS

A.J.C. and N.G.F. conceptualization; A.J.C. completed experiments; A.J.C. formal analysis; A.J.C. writing—original draft; A.J.C. and N.G.F. writing—review & editing; N.G.F. Supervision, funding support.

