## Supplementary Information for "Comparative analysis of wavelength-specific UV stress granule formation"

This Supplementary Information consists of Supplementary Methods, 7 Supplementary Figures, and Supplementary References.

**Supplementary Methods**

UV Meter Calibration

The UVA, UVB, and UVC sources were measured using a calibrated UV meter (THOR Labs, PM100D) and a UV sensor (THOR Labs, S120VC 200-1100nm). To ensure our UV meter and sensor were calibrated and reading accurately we used a known laser source at a known power and compared our UV meter readings to an identical UV meter that was previously calibrated by the company Light Conversion. This control UV meter belongs to the Titova Laboratory in the Physics department at Worcester Polytechnic Institute. Using the THOR Labs responsivity raw data (Supplementary Fig. S1) for our UV sensor we confirmed that both meters were outputting the same responsivity (mA/W) by measuring at least 3 different frequencies and at least 3 powers.^1^ The laser source used was a CARBIDE laser from Light Conversion (CARBIDE-CB3-40W) which is a 40W, 100kHz, 1030nm solid state laser with a pulse duration of 229 fs. This laser was used to pump OPA (optical parametric amplifier) with the continuous frequency range from 350nm/2.5um, pulse duration will slightly vary depending on the frequency. For measured frequencies the output was ~90fs. For each UV experiment, our UVA and UVB sources were allowed to heat up for at least 5 minutes and then using our UV meter and sensor the power was measured (mW). The UV radiation time required to reach a desired fluence (mJ/cm^2^) was then calculated by dividing the power (mW) by the area of the probe (0.71cm^2^).

Western Blotting

To verify that UV treatments induced cellular damage, we performed western blot analyses to assess damage-associated changes in protein phosphorylation. Phosphorylation of Erk1/2 (p-Erk1/2) has been previously reported in response to different wavelengths of UV exposure^2–4^. In addition to MAPK activation, poly(ADP-ribose) polymerase-1 (PARP-1) undergoes proteolytic cleavage following UV-induced stress, converting the full-length 116-kDa protein into two fragments of approximately 89 and 24 kDa^5,6^.

Protein extracts were prepared by directly lysing cells in 1x SDS sample buffer (62.5 mM Tris-HCl, pH 6.8, 4% glycerol, and 1.6% SDS). Lysates were boiled for 10 min at 95 °C and mechanically sheared by passing through a 22-gauge needle fitted to a 3-mL syringe at least 20 times. Dithiothreitol (DTT) was then added to each sample, and proteins were resolved on commercially prepared 12% Tris–glycine polyacrylamide gels (Thermo Scientific™) using 1x Tris–glycine running buffer (12.5 mM Tris base, 75 mM glycine, and 0.5% SDS) for approximately 90 min at 120 V. Proteins were transferred to polyvinylidene difluoride (PVDF) membranes using 1x transfer buffer (12.5 mM Tris base, 75 mM glycine, and 20% methanol) for 90 min at 100 V, with ice packs placed in the transfer chamber to maintain the temperature at ~4 °C.

Membranes were blocked in 5% nonfat dry milk (Demoulas Super Market Inc., Tewksbury, MA) prepared in 1x PBS–Tween (1% Tween) for at least 1 h at room temperature or overnight at 4 °C. Membranes were incubated with primary antibodies diluted 1:1,000 in PBS–Tween overnight at 4 °C, washed three times with 1x PBS–Tween, and then incubated with HRP-conjugated secondary antibodies diluted 1:10,000 in 5% milk in PBS–Tween. Protein bands were detected using SuperSignal™ West Pico PLUS Chemiluminescent Substrate (Thermo Scientific™) and visualized using an Azure c600 imaging system.

Cell Viability

In addition to stress response pathway activation, cell viability is also decreased in response to UV irradiation^2,7,8^. To confirm that our UV treatments are reducing cell viability, we measured cell viability using the MTT assay. Briefly, the MTT assay is used to detect mitochondrial respiration activity and in live cells mitochondrial respiration is consistent. Therefore, mitochondrial respiration is linearly related to the total amount of viable cells and can therefore be used to estimate cell viability^9^.

Cells were plated at 3,000 cells/well into 96-well plates. The following day cells were stress treated as indicated followed by a 2-hour incubation with MTT reagent (0.5mg/mL, Sigma) at 37ºC. MTT containing media was removed from wells and 100uL of DMSO was added to each well. Plates were placed into a shaker and incubated at 37ºC for 15 minutes. Absorbance for each well was read at 570nm.

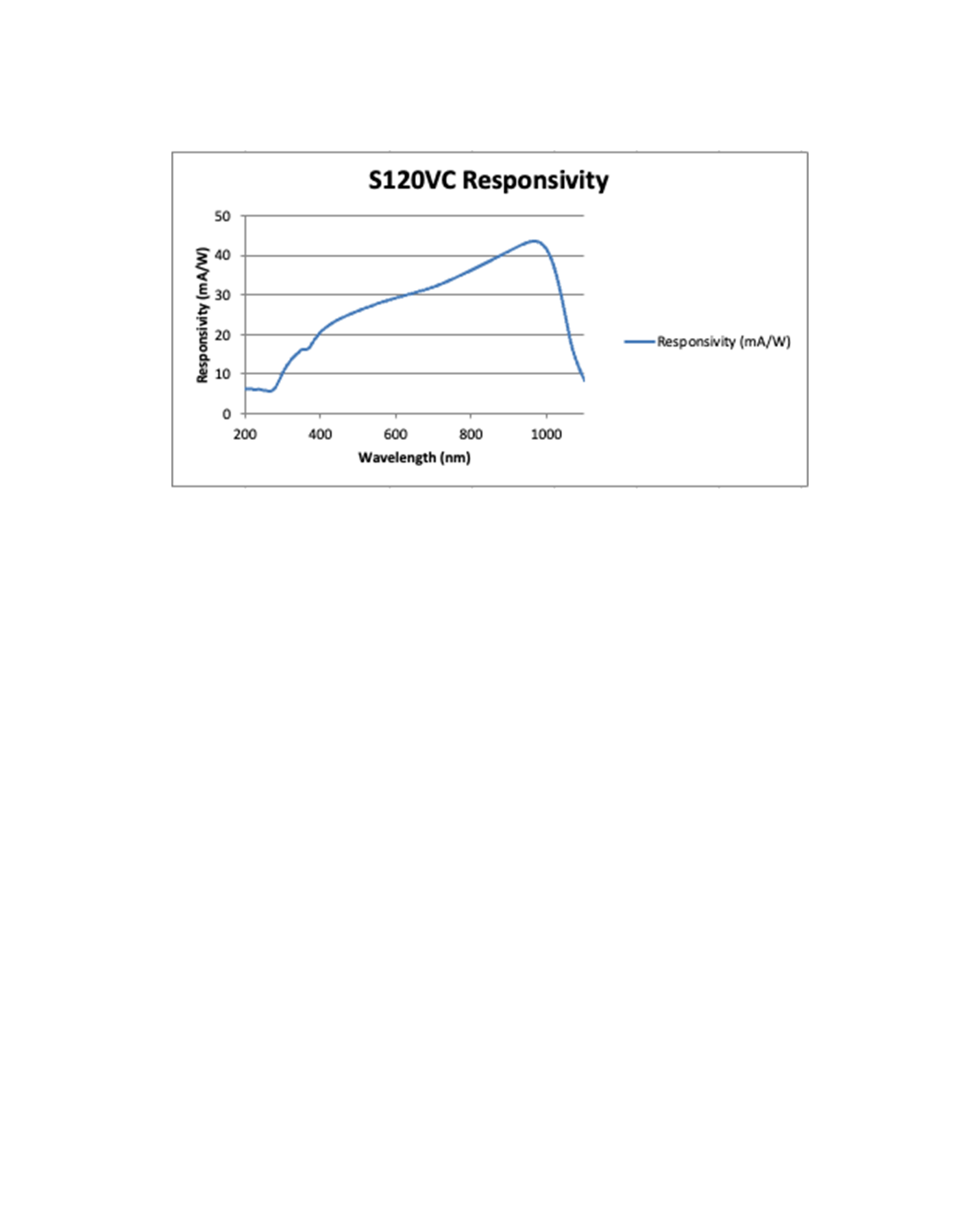

**Supplementary Figure S1.** THOR Labs ([THOR Labs responsivity raw data (S120VC)](https://www.thorlabs.com/newgrouppage9.cfm?objectgroup_id=3328)) raw data for responsivity of their S120VC (200-1100nm) UV sensor. This sensor was used throughout this study, and we ensured that it was measuring accurately according to this raw data.

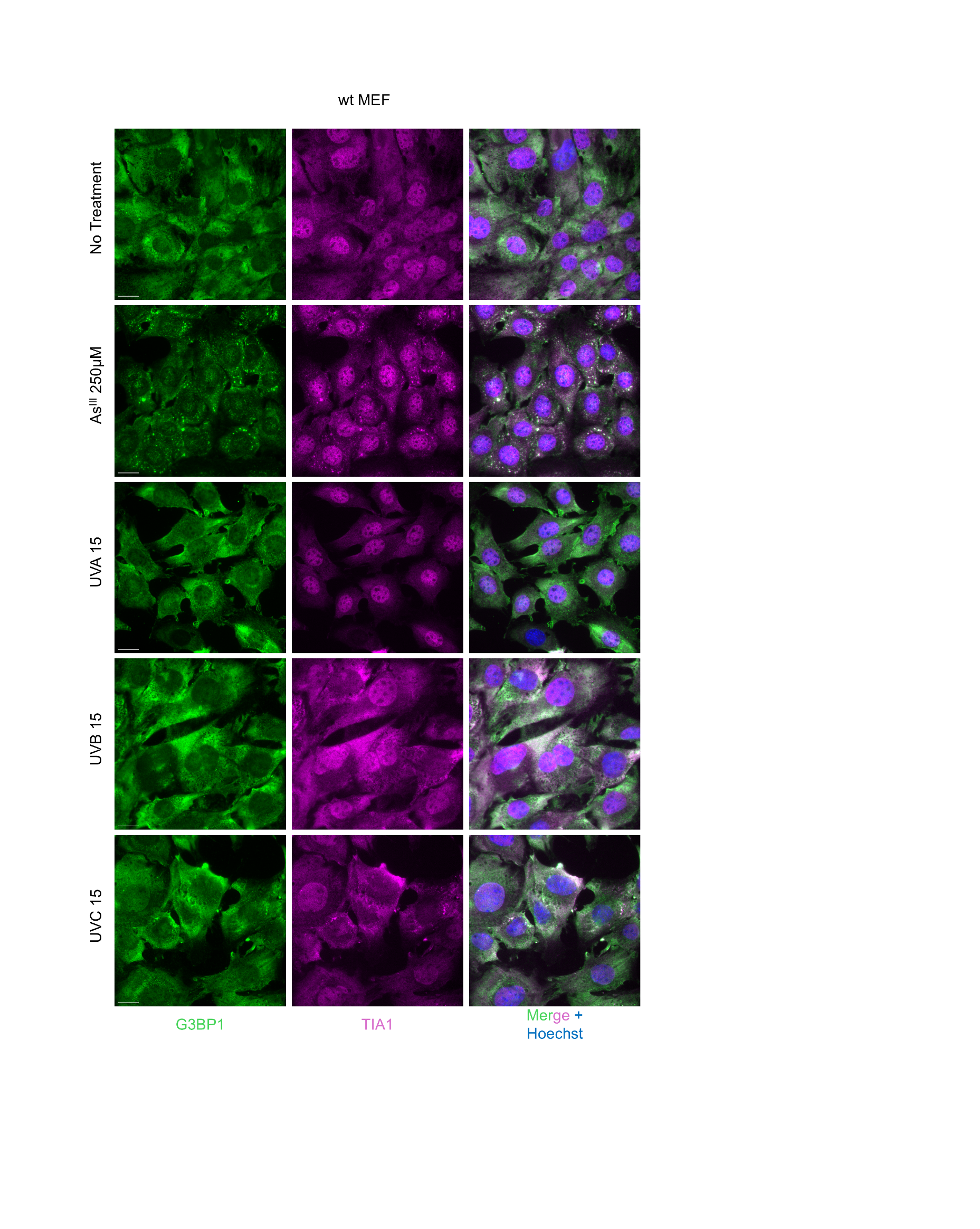

**Supplementary Figure S2. UVC induces minimal SGs in MEF cells.** Representative images of SG formation by G3BP1 and TIA1 signals in wt MEF cells treated as indicated. Cells were either untreated, treated with 250μM As^III^ for 1h, or exposed to 15mJ/cm^2^ of UV. UV samples were harvested and processed for imaging 4 hours after exposure. Scale bar = 20μm.

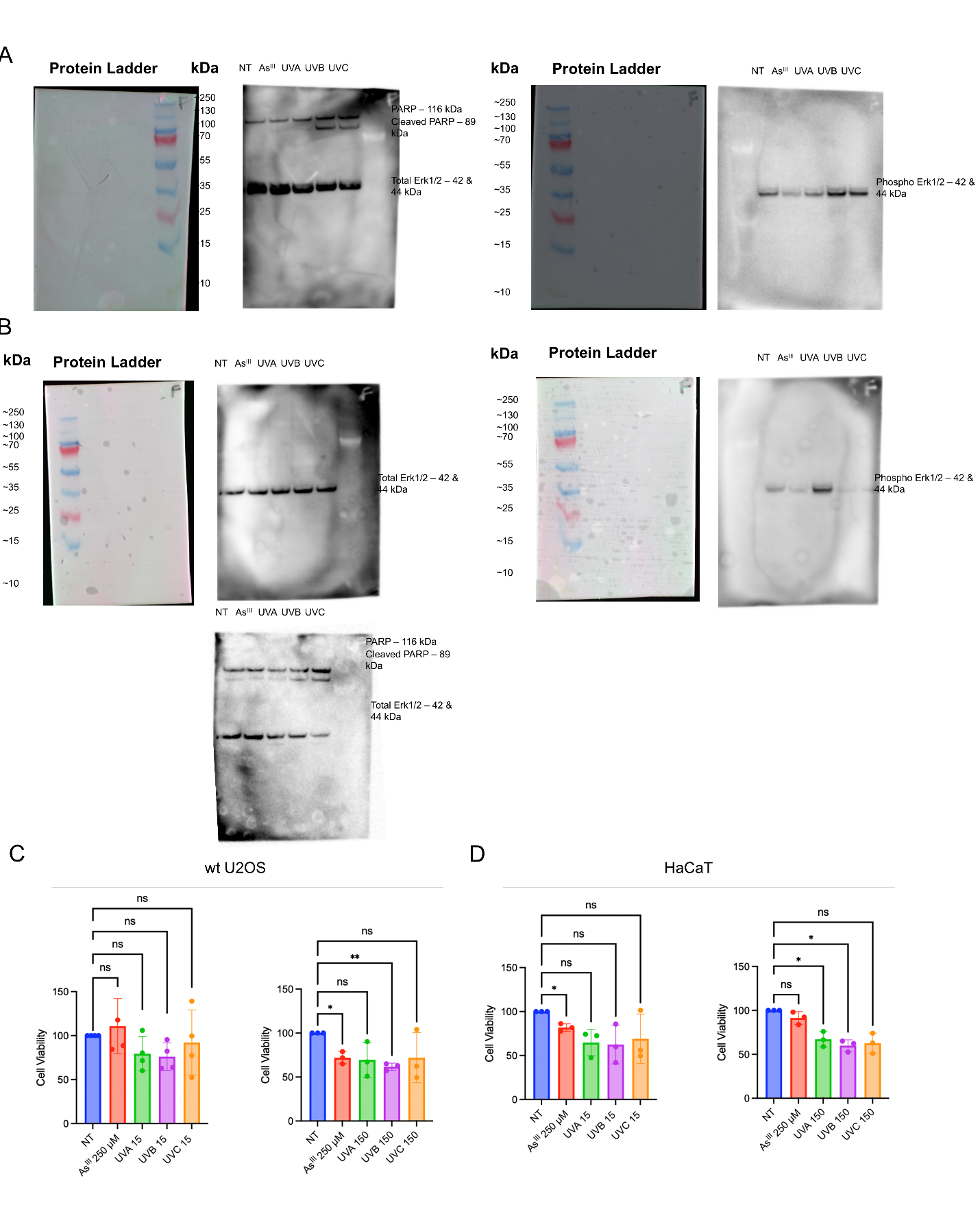

**Supplementary Figure S3. UVA, UVB, and UVC exposure induce cellular stress response pathways.** wt U2OS (A) and HaCaT (B) cells were left untreated, treated with 250μM As^III^ for 1h, or exposed to 150 mJ/cm^2^ of UVA, UVB, or UVC. UV samples were harvested 4 h post-exposure and processed for western lot analysis. Consistent with prior reports, our preliminary analyses revealed increased Erk1/2 phosphorylation and increased PARP cleavage following UV treatment. (C-D) Cell viability of wt U2OS (C) and HaCaT (D) cells after treated as indicated by MTT. Cells were either untreated, treated with 250μM As^III^ for 1h, or exposed to UV (15mJ/cm^2^ (left) or 150mJ/cm^2^ (right)). *n*=4 (C left panel) or *n*=3 (C right panel, D). We observed reduced viability under higher-dose UV conditions in both U2OS and HaCaT. Error bars are ±SD. *n*=3; error bars are ±SD; ns P > 0.05, ** P ≤ 0.01, **** P ≤ 0.0001 by one-way ANOVA and Dunnett’s post-hoc test.

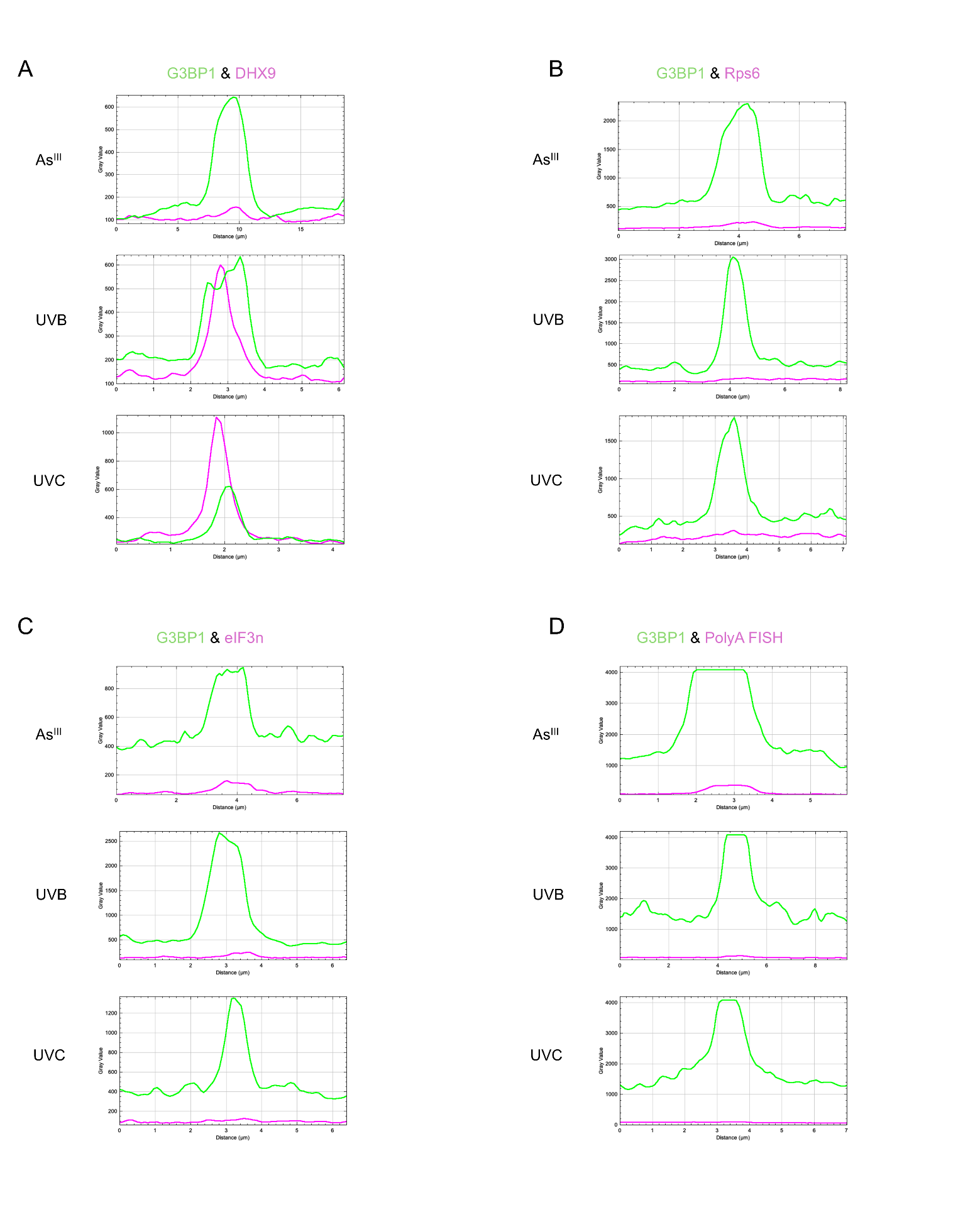

**Supplementary Figure S4.** **Co-localization plot profiles of stress granule–associated proteins in wt U2OS cells.** Plot profile analysis was used to assess co-localization of selected stress granule (SG)–associated proteins with G3BP1 under the indicated stress conditions. U2OS cells were left untreated, treated with 250 μM As^III^ for 1 h, or exposed to 150 mJ/cm² of UVA, UVB, or UVC. UV-treated samples were harvested 4 h post-exposure and processed for immunofluorescence. Images were acquired and analyzed using ImageJ. For each condition, a line was drawn through a representative region of the cell containing an SG, when present (in Fig. 3), and corresponding plot profiles were generated using ImageJ.

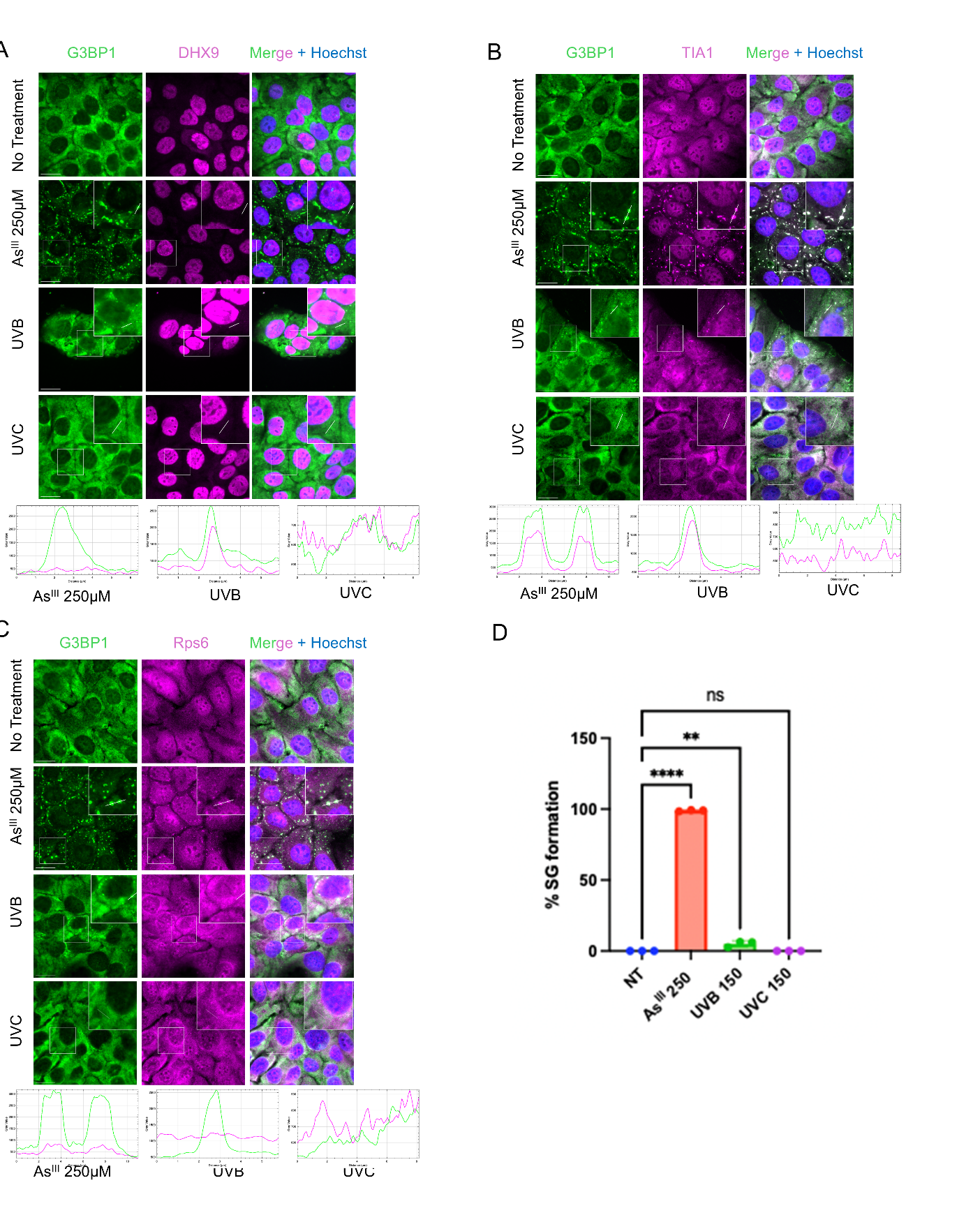

**Supplementary Figure S5.** **Stress granule quantification and co-localization analysis of stress granule–associated proteins in HaCaT cells.** Plot profile analysis was used to assess co-localization of (A) DHX9, (B) TIA1, and (C) Rps6 with G3BP1 under the indicated stress conditions. HaCaT cells were left untreated, treated with 250 μM As^III^ for 1 h, or exposed to 150 mJ/cm² of UVB, or UVC. UV-treated samples were harvested 4 h post-exposure and processed for immunofluorescence. Images were acquired and analyzed using ImageJ. For each condition, a line was drawn through a representative cellular region containing a stress granule (SG), when present, and corresponding plot profiles were generated to assess co-localization. In addition, the percentage of SG-positive cells (D) was determined by manually scoring at least 250 cells across three fields of view. *n*=3; error bars are ±SD; ns P > 0.05, ** P ≤ 0.01, **** P ≤ 0.0001 by one-way ANOVA and Dunnett’s post-hoc test. Scale bar = 20μm.

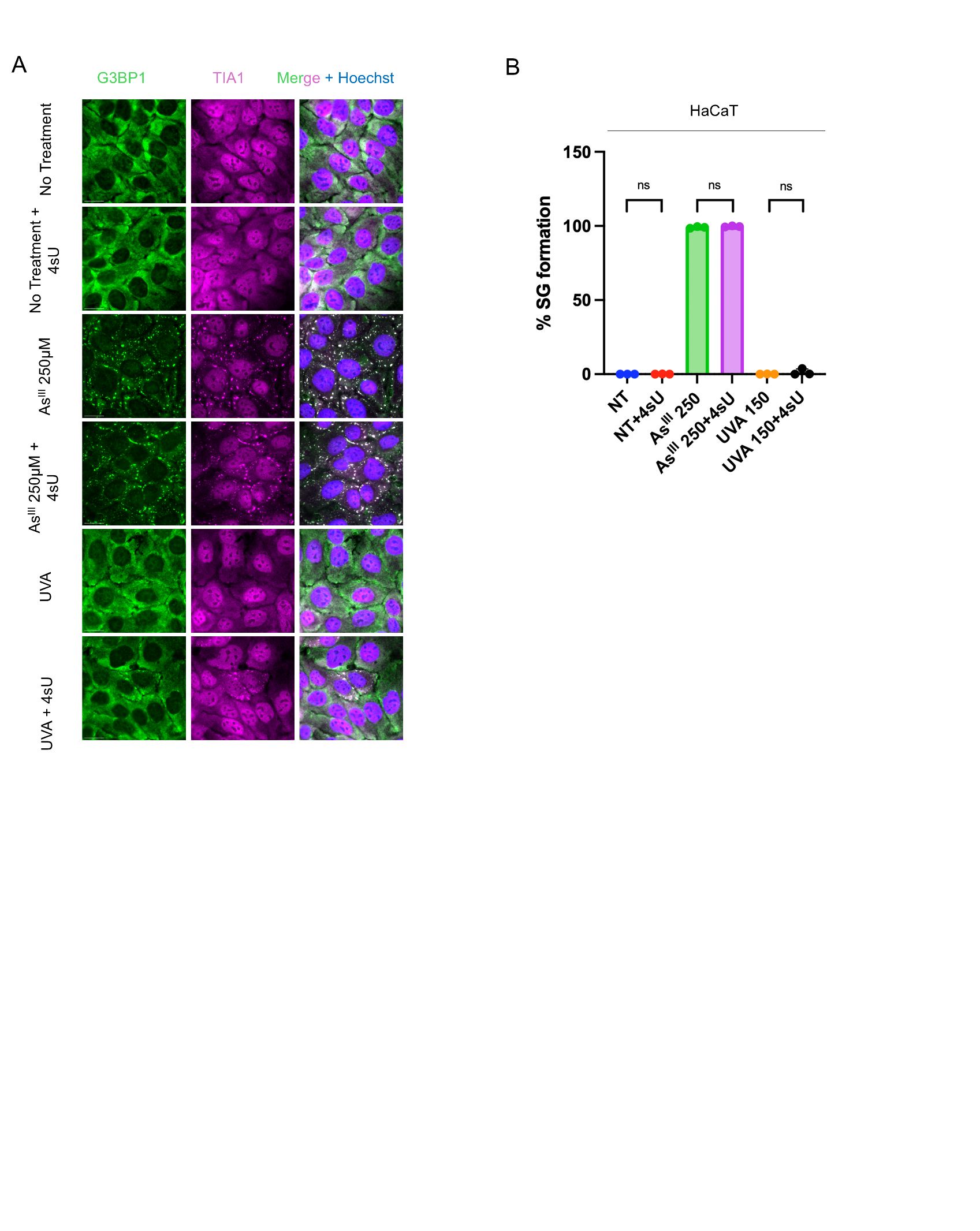

**Supplementary Figure S6. 4sU pretreatment does not robustly induce stress granule formation in HaCaT cells following UVA exposure.** HaCaT cells were left untreated or pretreated with 4sU (500ug/mL) for 1 h. Following pretreatment, cells were left untreated, treated with 250 μM As^III^ for 1 h, or exposed to 150 mJ/cm² of UVA. UV-treated samples were harvested 4 h post-exposure and processed for immunofluorescence. (A) Images were acquired and analyzed using ImageJ. (B) The percentage of SG-positive cells was determined by manually scoring at least 250 cells across three fields of view. *n*=3; error bars are ±SD; ns P > 0.05 by unpaired *t* test. Scale bar = 20μm.

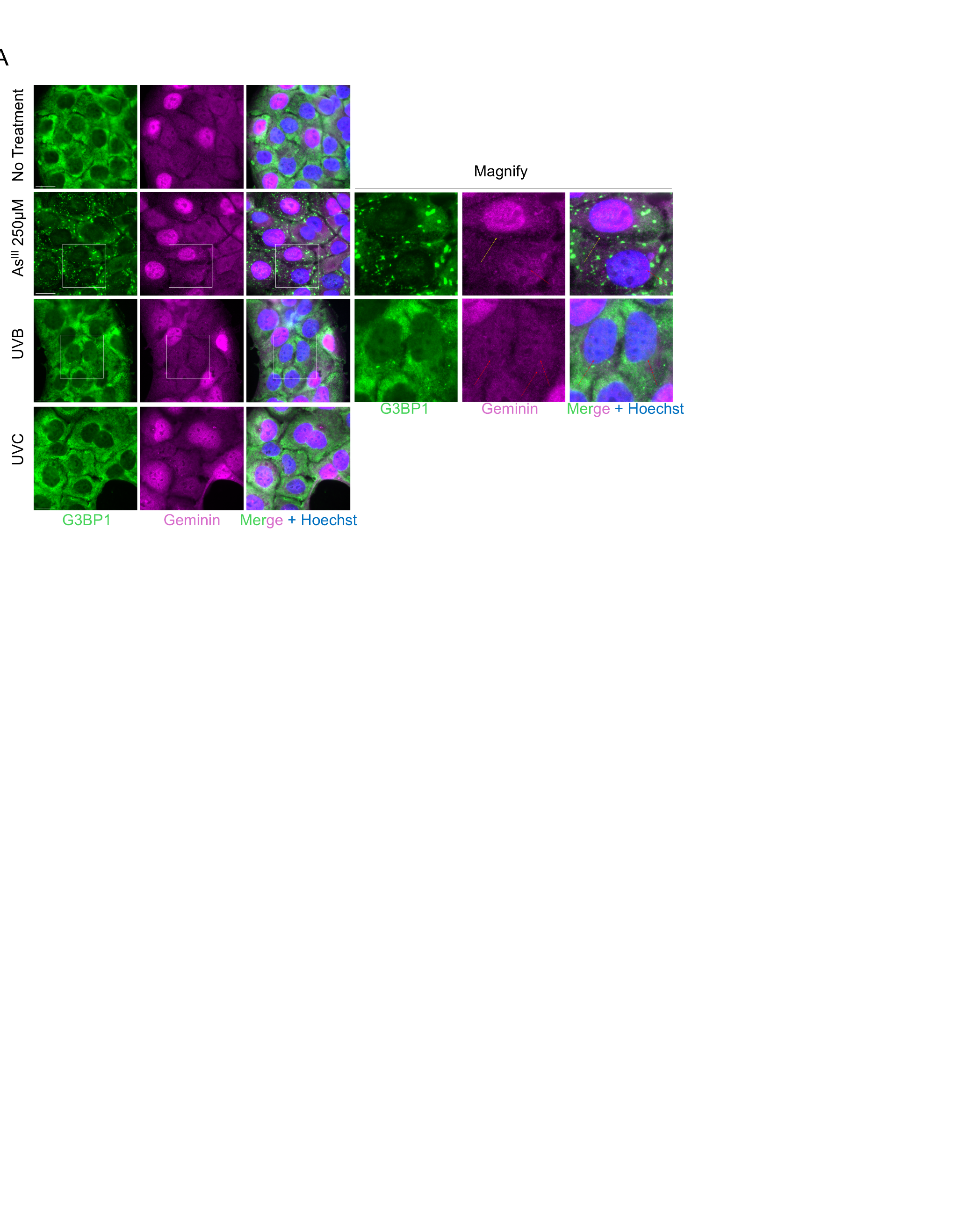

**Supplementary Figure S7.** **Nuclear geminin signal is absent in UVB-induced stress granule–containing HaCaT cells.** HaCaT cells were left untreated, treated with 250 μM AsIII for 1 h, or exposed to 150 mJ/cm² of UVA, UVB, or UVC. UV-treated samples were harvested 4 h post-exposure, stained for G3BP1 and geminin, and processed for immunofluorescence. Images were acquired and analyzed using ImageJ. Yellow arrows indicate cells in the S, G2, or M phases of the cell cycle, whereas red arrows indicate cells in the G1 phase. Scale bar = 20μm.
